# Intravital Two-Photon Imaging of Touch Sensory Axon Morphology in Mouse Skin

**DOI:** 10.1101/2025.09.08.674093

**Authors:** Sydney Kong, Emily Barnes, Gautham Chelliah, Shan Meltzer

**Affiliations:** Department of Pharmacology, Vanderbilt University

## Abstract

Low-threshold mechanoreceptors (LTMRs) are somatosensory neurons that detect innocuous light touch stimuli such as vibration, hair deflection, and pressure. They form subtype-specific terminals in the periphery and project axons centrally to the spinal cord to transmit tactile information. Current understanding about LTMR development and organization comes from fixed-tissue studies that cannot reveal the dynamic and temporal processes of neuronal wiring and remodeling. Here, we demonstrate a two-photon imaging method for visualizing LTMR axon morphology in the mouse right forepaw during development and in young adults. Two-photon microscopy can achieve high-resolution imaging within intact skin, allowing repeated imaging of the same axon terminals over postnatal timepoints. These approaches provide an *in vivo* system for the study of the cellular mechanisms that regulate LTMR patterning and plasticity. Its application to longitudinal analyses will make it possible to observe the assembly of touch circuits and their repair following injury. This technique may provide essential information about somatosensory axon structure and function in the skin.

**Summary:** This study presents an *in vivo* two-photon imaging approach to visualize the structures of low-threshold mechanoreceptor (LTMR) axon terminals in the forepaw skin in mice. By enabling repeated, high-resolution imaging of individual axons, the method provides a new platform for studying the sensory circuit structure and function during development and in adults.

## Introduction

Touch is the earliest sense to develop and plays a central role in how organisms experience and respond to the physical world. Touch is mediated by low-threshold mechanoreceptors (LTMRs), which are subtypes of dorsal root ganglion (DRG) neurons that detect innocuous mechanical stimuli, including light pressure, vibration, and hair movement ^1,2^. LTMRs are pseudounipolar neurons, and their peripheral axons innervate the skin to form subtype-specific terminals around structures. Their central axons relay tactile information to the central nervous system by forming synaptic connections with neurons in the spinal cord dorsal horn ^3^. However, how touch circuits are established and maintained is poorly understood.

Earlier foundational studies in cats during the 1960s and 1970s first identified these functionally distinct mechanoreceptors and their response properties in hairy skin ^4,5^. LTMR subtypes are classified by their unique morphology, conduction velocity, and response properties. The major subtypes found in the mouse hairy skin are Aβ rapidly adapting (RA) and slowly adapting (SA) LTMRs, Aδ-LTMRs, and C-LTMRs ^2,6,7^. Aβ RA-LTMRs, Aδ-LTMRs, and C-LTMRs form specialized endings called longitudinal lanceolate complexes around hair follicles ^2^. Axonal endings in the lanceolate complexes are closely aligned with hair shafts and are enveloped by finger-like processes from terminal Schwann cells, which provide trophic support and influence axonal morphogenesis ^8,9,10^. In glabrous skin, Aβ RA-LTMRs axons terminate in Meissner corpuscles, which are ovoid-shaped structures located within dermal papillae that are tuned to vibration and motion across the skin ^2,11^.

The spatial organization and density of LTMR terminals are influenced by both intrinsic genetic programs and skin-dependent cues during development ^12,13^. When damaged, LTMR dysfunction is associated with pathological conditions such as mechanical allodynia following nerve injury ^12,14^. Despite their importance in tactile function and pathological conditions, the *in vivo* mechanisms for LTMRs development and maintenance in the skin remain poorly understood. Much of what is known about their development stems from fixed-tissue analyses, which cannot capture the dynamic processes that may take place *in vivo* ^6,15–18^. For example, Aβ RA-LTMRs and Aδ-LTMRs exhibit active innervations around hair follicles and axonal pruning during early postnatal development ^19^. How hair follicle innervation and maintenance are achieved *in vivo* remains unknown.

Recent advances in two-photon laser-scanning microscopy, an advanced imaging technique that allows high-resolution visualization of fluorescently-labeled structures within living tissues, opened an exciting opportunity to capture the physiological dynamics of touch sensory axon endings in the skin. Two-photon imaging uses pulsed near-infrared light to excite fluorophores only at the focal plane, significantly avoiding photobleaching and phototoxicity and providing unprecedented capabilities for three-dimensional, spatially resolved structures within the tissue ^20^. Thus, biological structures within living tissues can be repeatedly imaged without damaging the cells themselves or surrounding tissues ^21,22^. Furthermore, the longer excitation wavelengths used in two-photon microscopy scatter less within tissue, allowing for deeper penetration and lower noise in complex environments ^23^. These advantages make two-photon microscopy well-suited for studying LTMR structures *in vivo*, especially where repeated, long-term imaging is necessary to track developmental and chronic changes over time.

Intravital imaging studies have revealed important insights into many physiological processes, from structural remodeling to neuronal activities. For example, two-photon imaging revealed structural synaptic plasticity and neuronal activities occurred in health and inflammation-induced remodeling of dorsal horn neurons in pain models ^24-26^. Protocols have been established for chronic *in vivo* two-photon imaging of hair follicles in the mouse ear skin ^21^. When paired with genetically modified mouse lines expressing subtype-specific fluorescent reporters, two-photon microscopy can also reveal individual neurons and their interactions with other neurons and non-neuronal cells in the skin microenvironment ^3^. In the somatosensory system, Thy1-YFP selectively labels Aβ LTMR axons, and Thy1-YFP labeled axons were positive for myelin basic protein and neurofilament, which are markers for myelinated Aβ axons ^27, 28^. Hence, Thy1-YFP is an excellent reporter for visualizing Aβ LTMR axons and their lanceolate complexes in the hair skin, and Meissner corpuscles in the glabrous skin. By contrast, *TrkB*^*CreER*^ selectively labels Aδ-LTMRs early during development, which form lanceolate endings around hair follicles but in a polarized pattern concentrated on the caudal side, a feature essential for their direction-selective responses to hair deflection ^7^.

This protocol explains the procedures for using two-photon microscopy to observe the structures of LTMR axons in the hairy and glabrous skins in developing and young adult mice. Our protocol can visualize axonal terminals with high resolution. The protocol includes several tips for optimal high-resolution imaging conditions and chronic imaging. This integrative approach allowed the investigation of the cellular mechanisms driving the assembly and plasticity of the touch sensory axons *in vivo*.

## Protocol

All experimental procedures were performed in compliance with institutional animal care regulations and were approved by the Institutional Animal Care and Use Committee (IACUC) at Vanderbilt University. Both male and female mice were included in the study. For the safety of the experimenters, all the procedures were performed with lab coats, sterile gloves, and masks. All tools were sterilized thoroughly prior to use, and all experimental surfaces were cleaned with 70% ethanol before and after each experiment.

1. Preparation of instruments
  1.1. Turn on the laser. Once on, the laser will take approximately 20 minutes to warm up.
  1.2. Turn on the two-photon microscope.
  1.3. Turn on heating pad and set it to 37°C to prevent hypothermia during anesthesia.
2. Preparation of anesthesia and the forepaw for imaging
  2.1. To induce the anesthesia, place the mouse in the induction chamber and set the system to 3.5% isoflurane until the mouse loses its righting reflex and demonstrates smooth, regular respirations.
  2.2. Place the mouse on the heating pad on the imaging platform and reduce the isoflurane to 1.5%. For postnatal mice, isoflurane is set to 2-4% to maintain anesthesia. Isoflurane should be delivered through the nose cone. Monitor anesthetic depth every 10 minutes by checking for loss of the pedal and palpebral reflexes observing respiratory rate and depth, mucous membrane color, and overall muscle tone.
  2.3. To reduce movement artifacts during imaging collection, including movement caused by breathing, the mouse should be placed on a different stage from the paw. For this protocol, the mouse body rests on a platform directly adjacent, but not touching, the microscope platform where the paw is mounted.
  2.4. Apply ophthalmic ointment to the eyes to prevent dryness during imaging.
  2.5. Gently apply depilatory cream to remove the hair on the paw using a cotton swab. After letting it sit for a minute, remove the cream and clean the mouse paw with 70% ethanol and water.
  2.6. Using a 30-gauge needle and black ink, tattoo two adjacent dots using gentle force on either side of the intended image region. Ensure that the dots are far enough away from the imaging field.
  2.7. On a glass microscope slide, place four dots of vacuum grease on the corners and place a small piece of clay on the middle of the slide. Place the slide under the objective lens and center the mouse’s paw on the slide, directly on the piece of clay **(Figure 1)**. Gently roll the mouse’s paw on the slide to flatten it and press it into the clay. Carefully place a coverslip on top and press down so the slide is secured. Adjust the angle of the mouse skin by putting pressure on each of the corners of the coverslip, until the surface of the coverslip and skin is parallel to the imaging plane.
  2.8. For imaging the glabrous skin, place the slide under the objective lens and center the mouse’s paw on the slide as detailed in 2.7. Use tweezers to rotate the paw upside down so that the glabrous skin faces up. Use the flat end of the tweezer to gently press the paw into the clay, trying to ensure the paw is as flat as possible. Once the paw is flat, place the coverslip on top and press down as described in 2.7.
3. Two-photon microscope imaging of LTMR morphology NOTE: The following parameters can be adjusted based on the specific experiment. Steps 3.1-3.6 describe the specific parameters used for the Thy1-YFP mice right forepaw, which can be used as a reference guide.
  3.1. To use a water-immersion objective lens, place a drop of distilled water on-top of the coverslip directly over the mouse paw.
  3.2. Carefully lower the objective lens until a water column is formed.
  3.3. Once the objective lens is centered over the mouse paw, close the microscope box and use the LED screen to focus on the sample. The tattoos on the mouse paw are visible on the LED screen as black spots. Using these landmarks, find and focus on the area in-between the black spots. This ensures that the same area of the mouse paw is imaged repeatedly.
  3.4. Once the sample is found, turn off the LED screen and switch the microscope to the “dichroic” setting.
  3.5. Turn on the tunable laser set at 960 nm and start scanning.
  3.6. Use the X, Y, and Z controllers, and adjust the zoom as needed, to find the desired area and start image acquisition software. A high-resolution image has a slice thickness of 1µm or 2µm.
  3.7. The imaging session will be limited to 25 minutes to minimize the duration of isoflurane exposure and reduce the risk of anesthesia-related adverse effects.
4. Postoperative care
  4.1. Remove the mouse paw from the glass slide, return the mouse to a recovery cage.
  4.2. Keep the mouse cage warm on the heating pad until the mouse recovers from the anesthesia.
  4.3. Place the mouse back into its home cage, observe the mouse for an additional 30 minutes to ensure the mouse integrates with its cage mates.
5. *In vivo* chronic imaging of LTMR morphology
  5.1. Place the mouse under the two-photon microscope by following steps 2.4 to 3.3.
  5.2. Using the LED screen, locate the ink dots from the previous imaging session. Orient the screen between the dots to ensure that the desired area is captured.
  5.3. Once the imaging area is found, follow steps 3.4 to 3.6 to continue performing *in vivo* imaging.
  5.4. Check the quality and intensity of images, as well as the morphology of major axons, obtained by two-photon excitation of the skin area. Ensure that the images fields are comparable to those taken during the previous imaging session.

## Representative Results

We used two-photon microscopy to visualize Aβ RA-LTMRs and Aδ-LTMRs axonal structures in the skin in young adults and during development. Mature lanceolate endings, that are fully developed, can be seen in young adult mice (∼postnatal day 20 and older) ^19,29^. We observed the overall organization axon terminals in the hairy skin in young adult mice, as shown in the maximum intensity projection image (**Figure 2A)**. The depth and shape of these axon terminals can be viewed using a three-dimensional reconstruction (**Movie 1**). Within this imaging field, hair follicles can be identified by their autofluorescence (**Figure 2A**). Lanceolate endings can be identified around guard and awl/auchene hair follicles and imaged further at high-resolution (**Figure 2C**). The maximum intensity projection of a mature lanceolate ending shows individual Aβ-LTMR axons encircling a hair follicle (**Figure 2C**), and the individual lanceolate ending can be visualized in single z-planes (**Figure 2D**). In the finger hairy skin, we observed immature lanceolate endings at P10, with fewer Aβ-LTMR lanceolate axons (**Figure 2E**). Individual z-planes illustrate that nascent lanceolate endings are sparse and lacks the distinctive elongated shape (**Figure 2F**).

**Figure 1:**
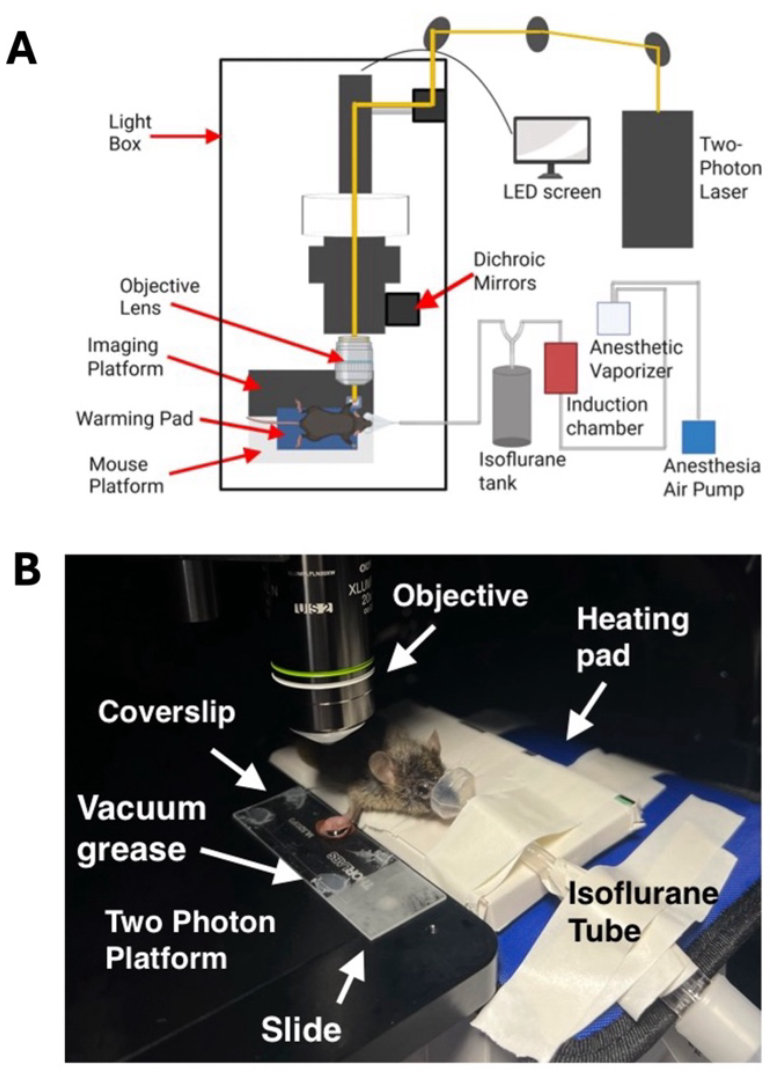
Schematic diagrams of the two-photon setup. (**A**) Schematic of the overall microscope and laser set-up. (**B**) Photo showing the mouse anesthetized on the stage and the mounted paw before imaging.

**Figure 2:**
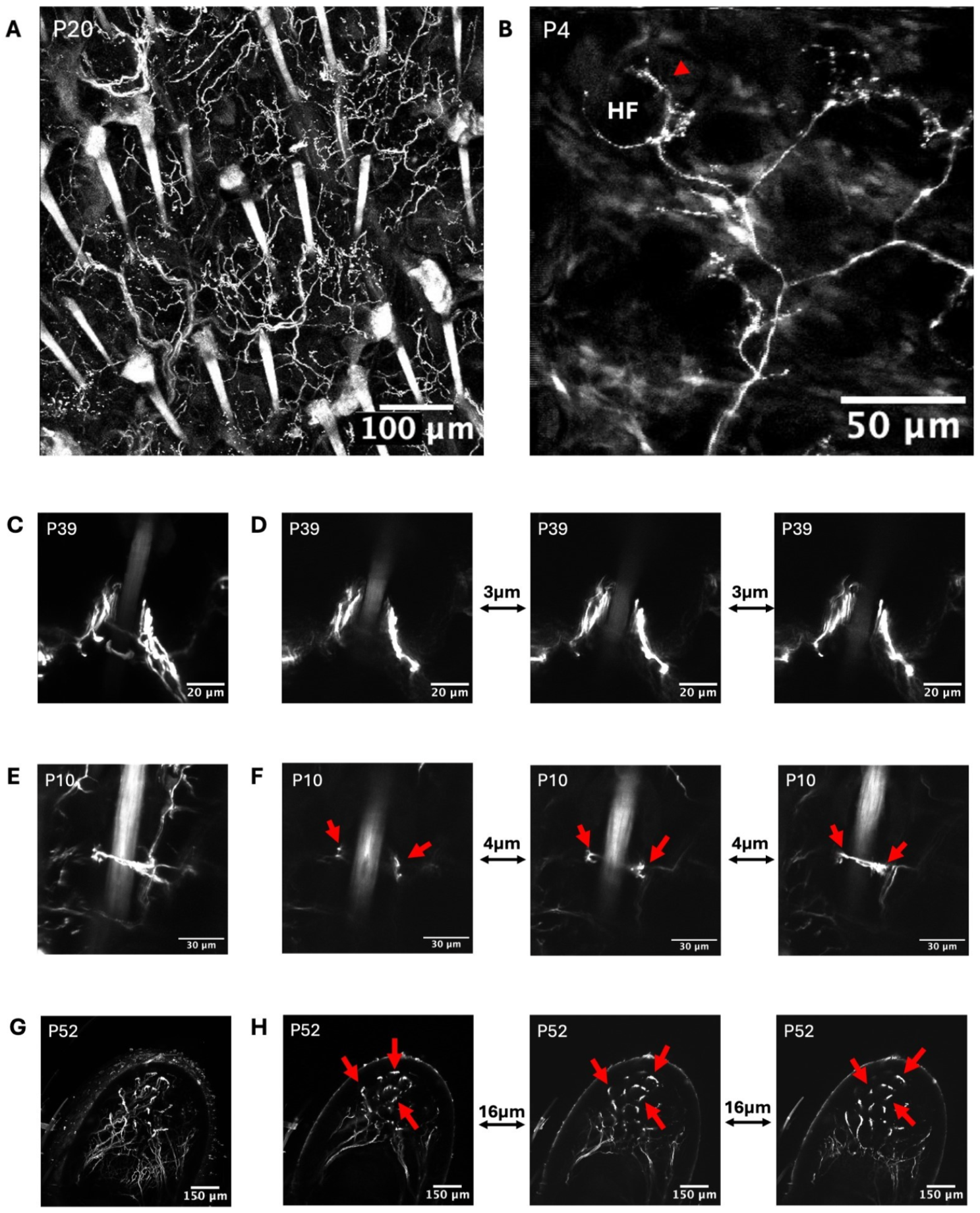
*In vivo* imaging of axon terminals in Thy1-YFP young adult mice. (**A**) Maximum intensity projection of a hairy skin image stack with 1 µm thickness taken from the hairy skin in the right forepaw. (**B**) Maximum intensity projection of a sparsely labeled Aδ-LTMR in the forepaw hairy skin at postnatal day 4. HF, hair follicle. Red arrowhead points to a hair follicle. Aδ-LTMRs are selectively labeled by *TrkB*^*CreER*^ driving the expression of a *Ai140* fluorescent reporter. Pregnant females were treated with 0.3 mg of tamoxifen at E12.5 to ensure specific labeling of Aδ-LTMRs. (**C)** Maximum intensity projection of a longitudinal lanceolate complex. (**D**) Selected focal planes showing individual lanceolate endings at a high resolution, each separated by 3 microns in z-planes. (**E**) Maximum intensity projection of an immature longitudinal lanceolate complex. **(F**) Selected focal planes showing the axonal progression of an immature lanceolate ending at high resolution, each separated by 4 microns. Arrowheads point out the developing Aβ RA-LTMRs. **(G)** Maximum intensity projection image of Aβ RA-LTMRs in the Meissner corpuscles in a glabrous fingertip in the Thy1-YFP mouse. **(H)** Individual frames taken from a glabrous fingertip image stack acquired at 4 µm z-steps, showing Aβ RA-LTMRs endings in Meissner corpuscles. Arrowheads point out Aβ RA-LTMRs in individual Meissner corpuscles. N ≥ 3 animals per condition.

Aδ-LTMRs are characterized by lightly myelinated somatosensory axons and can be selectively labeled by the combination of *TrkB*^*CreER*^ and Ai140 reporter during development ^7^. Pregnant females were treated with 0.3 mg tamoxifen at embryonic day 12.5 to ensure selectively labeling of Aδ-LTMRs. In the hairy skin, Aδ-LTMRs form lanceolate endings exclusively around non-guard hairs, with innervation and pruning occurring postnatally ^19^. By postnatal day 5, Aδ-LTMR axons form lanceolate endings around non-guard hairs follicles ^19^. A maximum intensity projection of mouse hairy skin illustrates one of these nascent crescent-shaped endings innervating a hair follicle at postnatal day 4 (**Figure 2B**). Of note, the pigmentation and autofluorescence from the surface of the developing skin was lower compared to adult mice, making this developing skin an ideal system for investigating somatosensory neuron development.

In the glabrous skin, Thy1-YFP selectively labels Aβ-LTMR axons that form Meissner corpuscles embedded in dermal papillae. To reduce background skin autofluorescence during imaging, the mouse’s paw was flattened against the coverslip, and the surface of the skin was cleaned with ethanol and water. Meissner corpuscles are ellipsoid in shape, consisting of stacked lamellar cells linked with sensory axon terminals ^2^. Their unique ultrastructure and superficial position within the epidermis organization allowed them to be highly sensitive to indentation and vibration ^9^. A maximum intensity projection image illustrates Aβ RA-LTMR innervation of Meissner corpuscles in the fingertip in adults (**Figure 2G**). Individual optical sections highlight discrete axon terminals within each corpuscle (**Figure 2H**). In these views, Meissner corpuscles appeared as elongated ellipsoids located just beneath the epidermis, and three-dimensional reconstruction show their axon terminals branching through stacks of lamellar cells (**Movie 2**), consistent with prior anatomical observations ^9,30^.

This technique can consistently image the same area of hairy skin over multiple days (**Figure 3**). Within the imaging field, individual axonal bundles can serve as landmarks. Some axons cluster together, forming structures that appear both thicker and brighter than the surrounding region, as indicated by the red arrows (**Figure 3**). While these structures can be traced throughout the images, it is important to note that the maximum intensity projections may not always be identical, likely due to slight set-up differences in paw positioning, flatness, and coverslip contact area (**Figure 3**).

**Figure 3:**
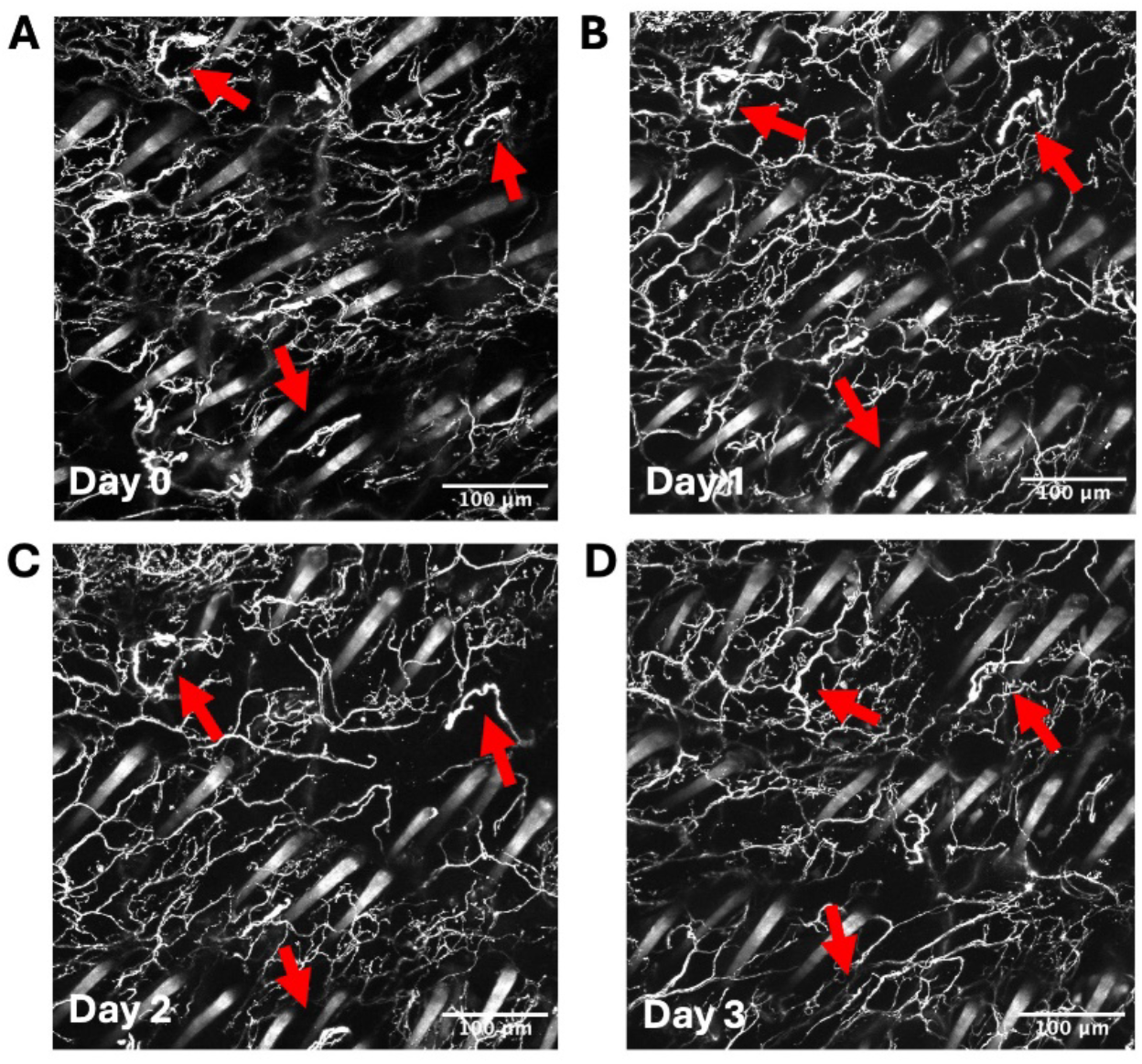
*In* vivo chronic imaging of axon terminals in Thy1-YFP young adult mice. The red arrows point to the individual axonal structures used to confirm that the same region is imaged. Maximum intensity projection of a hairy skin image stack (1 µm optical section thickness) acquired from the dorsal surface of the right forepaw of a mouse on **(A)** day zero, **(B)** day one, **(C)** day two, and **(D)** day three of imaging. N = 5 mice.

Altogether, these findings not only validate the ability of two-photon microscopy to reveal LTMR structures in intact hairy and glabrous skin but also establishes a framework for studying how LTMR development, function, and potential remodeling in disease or injuries.

## Discussion

This study demonstrates the use of two-photon microscopy for examining the morphology of low-threshold mechanoreceptors (LTMRs) in the intact skin of neonatal and young adult mice. Conventional histological approaches have offered foundational knowledge about LTMR subtype organization and target innervation ^6,7,31,32^, but they are limited to fixed time points and cannot capture the temporal progression of axonal changes. Our longitudinal imaging approach overcomes this limitation and allows for the observation of real-time axonal structure in a living animal. The imaging depth and resolution provided by two-photon microscopy allowed us to visualize axonal terminals in the skin without the need for tissue clearing or dissection, preserving the native microenvironment. Incorporating Aδ-LTMRs into this framework expands the utility of two-photon imaging beyond a single LTMR subtype, highlighting its potential to show how distinct mechanoreceptor populations develop, remodel, and interact with non-neuronal cell types in the skin, including Schwann cells, which are known to influence axonal growth and stability ^19,33^.

Several factors should be considered to obtain optimal and high-resolution imaging of LTMRs in the skin. First, genetically modified mice that express fluorescence in sparsely labeled neurons, for instance Thy1-YFP, allows for reliable imaging of the same population of neurons over time. Second, using clay to secure the paw onto the slide was important for adjusting the orientation of the paw and ensure the surface of the paw is perpendicular to the objective. Third, placing the body of the mouse on a separate platform from the paw greatly reduced imaging artifacts caused by breathing.

For chronic imaging, a number of landmarks can be employed to ensure consistent imaging of the same region. While visualizing the paw on the LED screen, the tattoo is detected as a black ink deposit, acting as a primary landmark. Then, during imaging, individual finger anatomy serves as additional landmarks as each digit can be visualized and traced to its connection with the paw, providing an estimate of the desired imaging field. Finally, confirmation of the imaging field can be achieved by comparing specific axonal morphology.

Future directions of research directions based on this imaging technique may include developmental dynamics of axonal activity, growth and regeneration. Integrating calcium imaging reporters with structural labeling may allow for simultaneous assessment of LTMR activity and morphology *in vivo*. Combining this system with injury models will be critical for exploring how LTMR circuits respond to peripheral nerve damage, and whether their regenerative trajectories reiterate developmental programs or invoke distinct repair mechanisms. Prior work has demonstrated the power of high-resolution imaging and optical clearing in direct visualization of Wallerian degeneration and axon growth within intact nerves ^34^. These studies demonstrate the value of tracking structural changes in peripheral nerves over time, supporting the rationale for applying a similar longitudinal strategy *in vivo* to examine how LTMR circuits respond to injury and whether or not their regenerative pathways mirror developmental processes or diverge into customized repair programs.

In conclusion, this study establishes a high-resolution platform for imaging LTMR during development and in adults *in vivo*. By visualizing axon dynamics in their native context, new insights can be gained into the dynamic processes that shape the somatosensory system and lay the groundwork for future investigations into the mechanisms of touch perception and repair.

## Acknowledgements

We would like to thank Dr. Hannah Elam, Joel Rodriguez Troncoso, John Girbert, Dr. Snighda Mukerjee, Rachelle Larivee, and Dr. Tegy J. Vadakkan for all of their support and assistance throughout this research. We thank Ryan Michael Nuera for help with setting up the isoflurane system. This work is supported by Howard Hughes Medical Institute Hanna Gray Fellowship (Grant# GT17518).

Materials

Materials:
| Name | Company | Catalog Number | Notes |
| --- | --- | --- | --- |
| Anesthesia Air Pump (HF), 110V | RWD Life Science Inc. | R510-30 | Isoflurane System |
| Compact Small Animal Anesthesia Machine, Isoflurane, Easy Fill, 0-4L | RWD Life Science Inc. | TAIJI-IE | Isoflurane System |
| Anesthesia Induction Chamber – Mouse and Rat | RWD Life Science Inc. | V105 | Isoflurane System |
| Anesthesia Concentric Mask | RWD Life Science Inc. | R510-M | Isoflurane System |
| Gas Filter Canisters | RWD Life Science Inc. | R510-31-6 | Isoflurane System |
| Small Animal Anesthesia Operation Platform | RWD Life Science Inc. | R510-PGL | Isoflurane System |
| Cone Mask with Tubing for Neonatal Mice | RWD Life Science Inc. | 68680 | Isoflurane System |
| 20X Objective Lens – Super NA 1.0 | Olympus | 1-U2B965 | Two-Photon Microscope Setup |
| Isoflurane, USP (250 mL) | Piramal Critical Care | 6679-017-25 | Isoflurane System |
| High Vacuum Grease | Sigma Aldrich | Z273554 | Imaging System |
| Laser Cover Box | Teco Technology |  | Imaging System |
| Chameleon Discovery NX with TPC: Dual | Coherent | 1385777 | Two-Photon Microscope Setup |
| Bruker Ultima Investigator Plus Laser Scanning Microscope | Bruker Nano, Inc. | INVP-01 | Two-Photon Microscope Setup |
| Mice | Charles River |  |  |
| Rigid Stand with Large Rectangular Insert Holder | Thorlabs | MP15M | Two-Photon Microscope Setup |
| Far Infrared Warming Pad with Controller (AC power) | Kent Scientific | RT-0515 | Two-Photon Microscope Setup |
| Nair Hair Remover Cream | Amazon | 28001 | Two-Photon Microscope Setup |
| 70% Ethanol | iLab Vanderbilt |  | Two-Photon Microscope Setup |
| Soothe® Lubricant Eye Ointment | Bausch + Lomb |  | Two-Photon Microscope Setup |
| 22x60 Cover Slip | VWR | 16004-330 | Two-Photon Microscope Setup |
| SuperFrost Microscope Slides | Fisher Scientific | 12-550-15 | Two-Photon Microscope Setup |
| Air-Hardening Modeling Clay | DAS® |  | Two-Photon Microscope Setup |
| Butterfly Needles | Fisher Scientific | 1484041 | Two-Photon Microscope Setup |

